# A chromosome-level phased *Citrus sinensis* genome facilitates understanding Huanglongbing tolerance mechanisms at the allelic level in an irradiation-induced mutant

**DOI:** 10.1101/2022.02.05.479263

**Authors:** Bo Wu, Qibin Yu, Zhanao Deng, Yongping Duan, Feng Luo, Frederick Gmitter

## Abstract

Sweet orange (SWO) originated from introgressive hybridization of pummelo and mandarin resulting in a highly heterozygous genome. Here, we assembled a chromosome-level phased Valencia SWO (DVS) genome with ∼98.5% completeness, high accuracy (QV=50.6), and the highest annotation BUSCO completeness (99.2%) thus far in citrus. DVS harbors a high level of allelic variances and enables study of allelic somatic structural mutations and corresponding allelic expression alteration in two SWO mutants, one with high Huanglongbing tolerance (T19) and one more sensitive (T78). In T78, a large deletion on the pummelo-origin chr8 causes regional allelic expression absence. In T19, seven upregulated genes are located at one terminal of a translocated segment, including three genes related to heat shock protein (HSP) regulation. Furthermore, 68 of 133 *HSPs* are significantly upregulated in T19, which may be related to its enhanced HLB tolerance by preventing phloem necrosis. The DVS will advance allelic level studies in citrus.

## Introduction

Sweet orange (*Citrus sinensis* L., SWO) has more than 2,300 years of recorded domestication history^1,2^, and most of its cultivars could date back to a single plant through somatic mutations^3,4^. The first SWO originated from complex hybridization processes involving mandarins (*Citrus reticulata* Blanco) and pummelos [*Citrus maxima* (Burm.) Merr.]^2,3^. Several other citrus cultivar groups also arose from interspecific introgressive hybridization events^5^, such as lemons (*Citrus limon* L.) and grapefruit (*Citrus paradisi* Macfad.). The primitive *Citrus* species contributing to the hybridizations generally diverged between 3 and 8 million years ago and vary substantially in genomes and phenotypes^5,6^. Interspecific and introgression hybrids in citrus are phenotypically distinct from their parents. Meanwhile, diverse SWO cultivars from somatic mutations are valuable resources for horticultural trait studies^4,7^. A phased genome assembly with allelic information revealed is fundamental for understanding the special genetic characteristics of SWO. With the haploid SWO reference genome, the somatic mutation calling and gene expression quantification in these mutants could be compromised in the highly divergent allelic genome regions. Moreover, a high-quality phased SWO reference genome will facilitate allelic level analysis in interspecific or introgression hybrids between pummelos and mandarins.

Due to difficulty of assembling the highly heterozygous genome, the current best SWO reference genome is assembled from a di-haploid sweet orange (HSO)^2,4^. Later efforts to sequence diploid SWOs only generated similar haploid assembly sizes with quality inferior to HSO^3,4^. Although the mapping-based phasing method, which is designed based on genomes with intraspecific heterozygosity, partitioned the diploid SWO genome into 325 phased blocks, it failed to provide a phased reference-level genome^4^. Here, we assembled a chromosome-level phased Valencia sweet orange (DVS) genome with ∼98.5% completeness and high accuracy (QV=50.6). The gene structure annotation of DVS has the highest BUSCO completeness (99.2%) thus far in the genus, representing a ∼6.2% improvement compared to the previously published di-haploid sweet orange genome. DVS harbors a high level of allelic variance, significantly higher than previously reported from studies using mapping-based methods. Approximately 12.7% of sweet orange genes have an allele affected by disruptive small variant(s), and 28.2% of the genes have an allele affected by structural variant(s). In transcriptome sequencing data, the alleles of 49.6% ± 0.07% genes were expressed with ≥ 1.5-fold difference in sweet orange. DVS enables study of allelic somatic structural mutations and corresponding allelic expression alteration.

Moreover, the relative genetic uniformity of sweet orange cultivars and their use in monoculture production also makes them vulnerable to disease epidemics. Citrus Huanglongbing (HLB) is a devastating disease mainly caused by *Candidatus* Liberibacter asiaticus (*C*Las)^8^. All commercial SWO cultivars are susceptible to HLB, and the selection of HLB tolerant/resistant germplasm has been considered the ultimate solution. Although no absolute HLB immunity has been found in natural citrus germplasm, different degrees of HLB tolerance and sensitivity have been observed^9– 15^. However, no genetic variations have been correlated with the high HLB-tolerance. In this study, we have selected an irradiation-induced Valencia SWO mutant with long-lasting (over 15 years) HLB tolerance. By taking advantage of our nearly complete phased DVS genome, we were able to reveal the molecular mechanism underlying its high HLB tolerance at the allelic level.

## Results

### Phased Valencia SWO genome assembly

We obtained 143.9 Gb (∼ 420 ×) whole-genome PacBio continuous long reads (CLR) for an ordinary diploid Valencia SWO (DVS). The assembly process was optimized according to the allelic divergence in the SWO genome. We first obtained a 607.6 Mb raw assembly using CANU, including 383 contigs with an N50 length of 23.5 Mb (estimated using the haploid genome size) or 15.4 Mb (using the total assembly size) (Supplementary Table 1). Then we applied phased assembly on the collapsed and expanded regions (Extended Data Fig. 1A). Approximately 3.9 Mb collapsed regions in 14 regions remain unphased in the final assembly, with 3.2 Mb at the 5’ end of chr2 (Fig. 1A). By resolving the repetitive units of long tandem-repeats (Extended Data Fig. 1B), we connected all filtered contigs into 18 pseudo-chromosomes totaling 598.6 Mb, which were assigned into two homologous chromosome sets DVS_A (chr1-9A, primarily mandarin-origin) and DVS_B (chr1-9B, primarily pummelo-origin) (Fig. 1A). An average hamming error rate of 0.18% was observed across the genome, and no switch error was detected in the interspecific heterozygous regions with pummelo and mandarin reference.

**Fig. 1:**
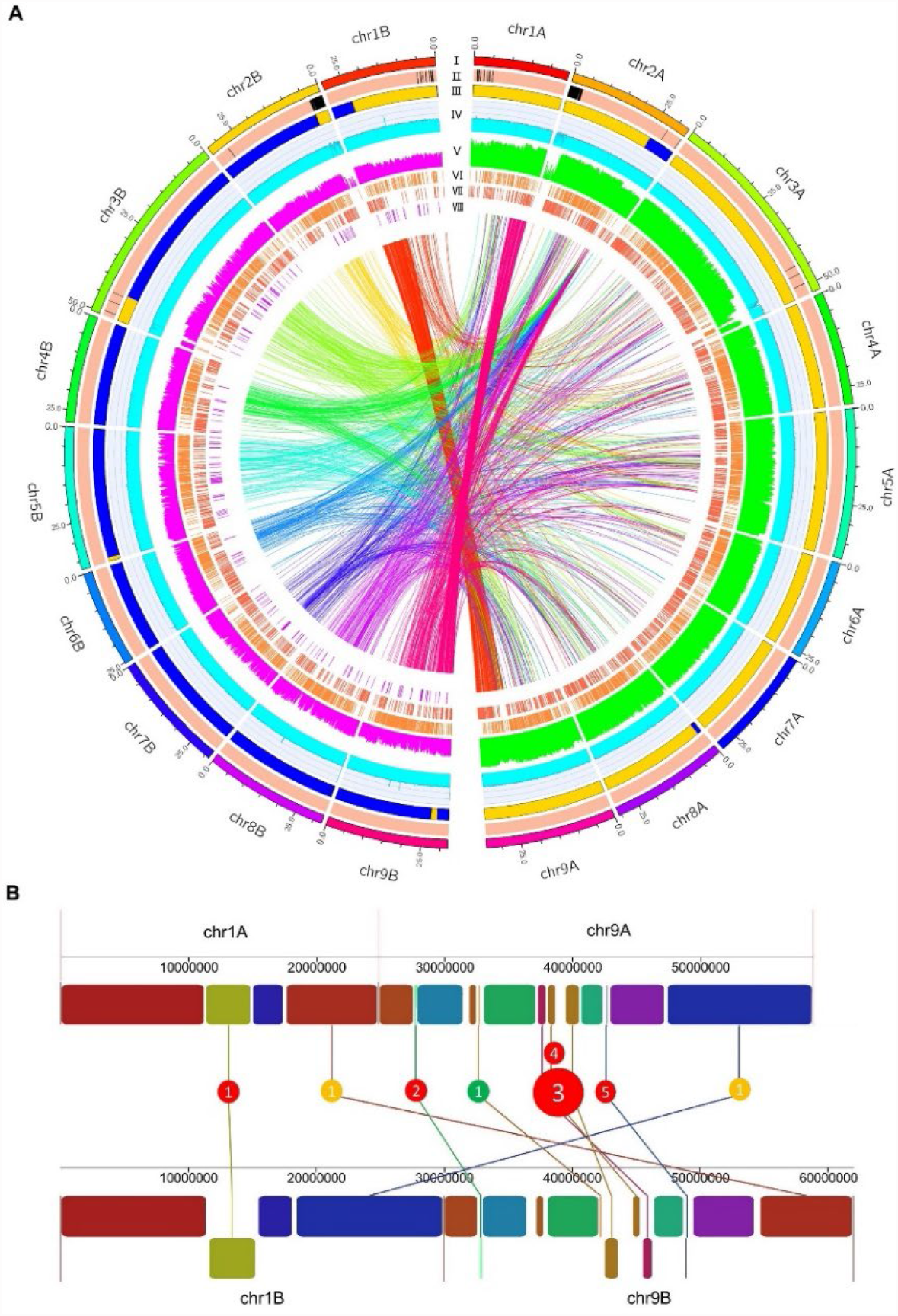
Characterization of the DVS genome and its intra-genomic variations. **A**. Characteristics of the phased DVS assembly and whole-genome variation distribution. (I) Ideogram of the DVS genome. Non-homologous chromosomes were colored distinctly, and homologous chromosomes were colored with different depths of the same colors. The unit of the tick labels is million base pairs (Mb). (II) The phased (light red) and unphased region (black) in the assembly. (III) Distribution of the mandarin (orange) and pummelo (blue) origin chromosomal regions. (IV) Sequencing coverage (binary logarithm shown in the graph) of uniquely mapped CLR reads across the assembly in 50 kb windows across the DVS. Neighboring windows were overlapped by 20 kb. The vertical axis ranged from 0 to 16. (V) Histogram of the binary logarithm of indel (purple) and SNV (green) counts in 100 kb non-overlapping continuous windows. The vertical axis ranged from 0 to 12. (VI) and (VII). Tandem duplications (DUPs, VIII) and insertions (INSs, VII) were detected by comparing DVS_A with DVS_B. These DUPs and INSs could also be described as deletions in the other chromosome set. (VIII) Inversions (INVs) in DVS_B with DVS_A as the reference. The inner links showed all identified inter-chromosomal translocations (TRA) by comparing DVS_A and DVS_B. The links started from each DVS_B chromosome have the same color as the chromosome in the ideogram. **B**. The seven largest SVs on chr1 and chr9. Rectangles with the same colors on chr1/9A and chr1/9B denote the orthologous blocks. The rectangles reversed in direction were shown in the bottom row on chr1B and chr9B. Red, yellow, and green circles denote INVs, inter-chromosomal TRAs, and intra-chromosomal TRAs, respectively.

The DVS assembly was estimated with 98.5% (K-mer) or 98.7% (BUSCO) completeness and an average error rate of 8.8E-6 (QV=50.6). Its base error rate is 46.8-fold lower, and its K-mer based completeness is 25.6% higher than HSO (if not specified, HSO denotes HSO v4) (Supplementary Table 2 and Extended Data Fig. 2A). DVS_A and DVS_B have a good syntenic relationship with HSO except for several structural variation (SV) affected regions (Extended Data Fig. 2B). With DVS as the reference, a 99.9% mapping rate was achieved on the DVS whole-genome NGS data. Higher mapping rates were observed on 12 tested SWO datasets with DVS as the reference compared to HSO (Extended Data Fig. 2C and Supplementary Note 1). DVS_A and DVS_B have similar BUSCO completeness scores (98.4% and 98.3%) with HSO (98.4%), but HSO is on average 37.3 Mb larger. The extra regions in HSO are mainly located in its arbitrarily connected pseudochromosome chrUn (Extended Data Fig. 2B) which has an error rate of 0.21%. Compared with DVS, HSO has higher proportions of low-coverage and long tandem-repeat regions (Extended Data Fig. 2D) that were filtered and connected through repeat-unit resolving in DVS.

### Chromosomal origin and intra-genomic variations of DVS

The origin of the DVS chromosomal regions was inferred with 20 pummelos (P) and 20 mandarins (M) (Supplementary Table 3). DVS_A contains ∼290.3 Mb (97.1%) M and ∼8.7 Mb (2.9%) P regions, and DVS_B has ∼37.7 Mb (12.6%) M and ∼261.9 Mb (87.4%) P regions (Fig. 1A and Supplementary Table 4). DVS contains approximately 254.8 Mb (84.7%) P/M, 37.5 Mb (12.5%) M/M, and 8.6 Mb (2.8%) P/P regions.

DVS_A and DVS_B share an 96.2% overall nucleotide similarity. 4,353,521 SNVs, 152,977 small indels (< 50 bp), and 9,989 SVs were detected between them (Fig. 1A). The SVs include 4,923 insertions (INSs), 4,563 deletions (DELs), 170 tandem duplications (DUPs), 156 inversions (INVs), and 177 translocations (TRAs). The seven largest SVs on chr1 and chr9, including two TRAs and five INVs, are shown in Fig. 1B. The inter-chromosomal recombination between chr1A and chr9A is shared by all three Valencia SWO accessions sequenced in this study.

### DVS gene structure annotation and allelic gene statistics

Approximately 49.0% (293.4 Mb) of the DVS genome were predicted to be transposable elements (Extended Data Fig. 3). A total of 55,745 protein-encoding genes were annotated, including 27,807 on DVS_A and 27,938 on DVS_B. The DVS annotation has 99.2% completeness by BUSCO test, 6.2% higher than HSO and the highest among published citrus genomes (Extended Data Fig. 4A). The protein sequences from DVS and six other citrus assemblies were phylogenetically clustered into 24,817 citrus orthogroups (COGs). The COGs were classified into 19,328 high-quality (HQ) and 5,489 low-quality (LQ) COGs based on their homology to plant proteins. DVS, DVS_A, and DVS_B have the most HQ-COGs and the fewest LQ-COGs (Extended Data Fig. 4B). A total of 1,386 HQ-COGs with members in DVS are absent in HSO, and 549 are the other way around (Supplementary Table 5). There are 533 HQ-COGs present in the two parental species genomes (HWB and MSYJ) but missing either in DVS_A or DVS_B (Extended Data Fig. 4C and Supplementary Table 5). Forty-seven high-quality COGs present in pummelo and mandarin are missing in both DVS and HSO (Supplementary Table 5).

DVS owns 22,614 orthologous (allelic) gene groups (DOGs), including 17,693 containing colinear orthologous gene pairs between DVS_A and DVS_B (Supplementary Table 6). The allelic genes from DVS_A and DVS_B share an average SNV density of 15.3 / kb in the exonic regions. The orthologs of 12.7% (7,069) genes are affected by 2.0 ± 1.0 high impact (disruptive such as frameshifting) intra-genomic small variants. 28.2% (15,721) genes are overlapped with at least one intra-genomic SVs. 3,408 (12.2%) DVS_A genes and 3,452 (12.4%) DVS_B genes are monoallelic without allelic genes found in the other chromosome set (Supplementary Table 7). These monoallelic genes are significantly (FDR < 0.05) overrepresented in gene ontologies including defense response, sexual production, and hormone signaling (Extended Data Fig. 4D and Supplementary Table 8).

### Irradiation-induced Valencia sweet orange mutants

We took advantage of the allelic information of the DVS assembly to probe the possible underlying molecular mechanisms in an HLB-tolerant SWO mutant. Most trees growing in the same field trial location as the mutants were killed by HLB or removed because of severe decline caused by HLB. Only six trees from two original selections, four from T19 and two from T78, are still growing in the grove. The four T19 trees included one with a lost tag (SF), which was proven identical to the other three T19 trees by whole-genome sequencing, as described in the following section. Though similar *C*Las titers were detected on T19, T78, and DVS (Fig. 2A) indicating equivalent infections, the T19 trees had significantly greater (p < 0.01) leaf area indexes (LAIs) than DVS and T78 (Fig. 2B). The four T19 trees are still healthy (Fig. 2C), while severe symptoms have developed on DVS (Fig. 2D) and the two T78 trees.

**Fig. 2:**
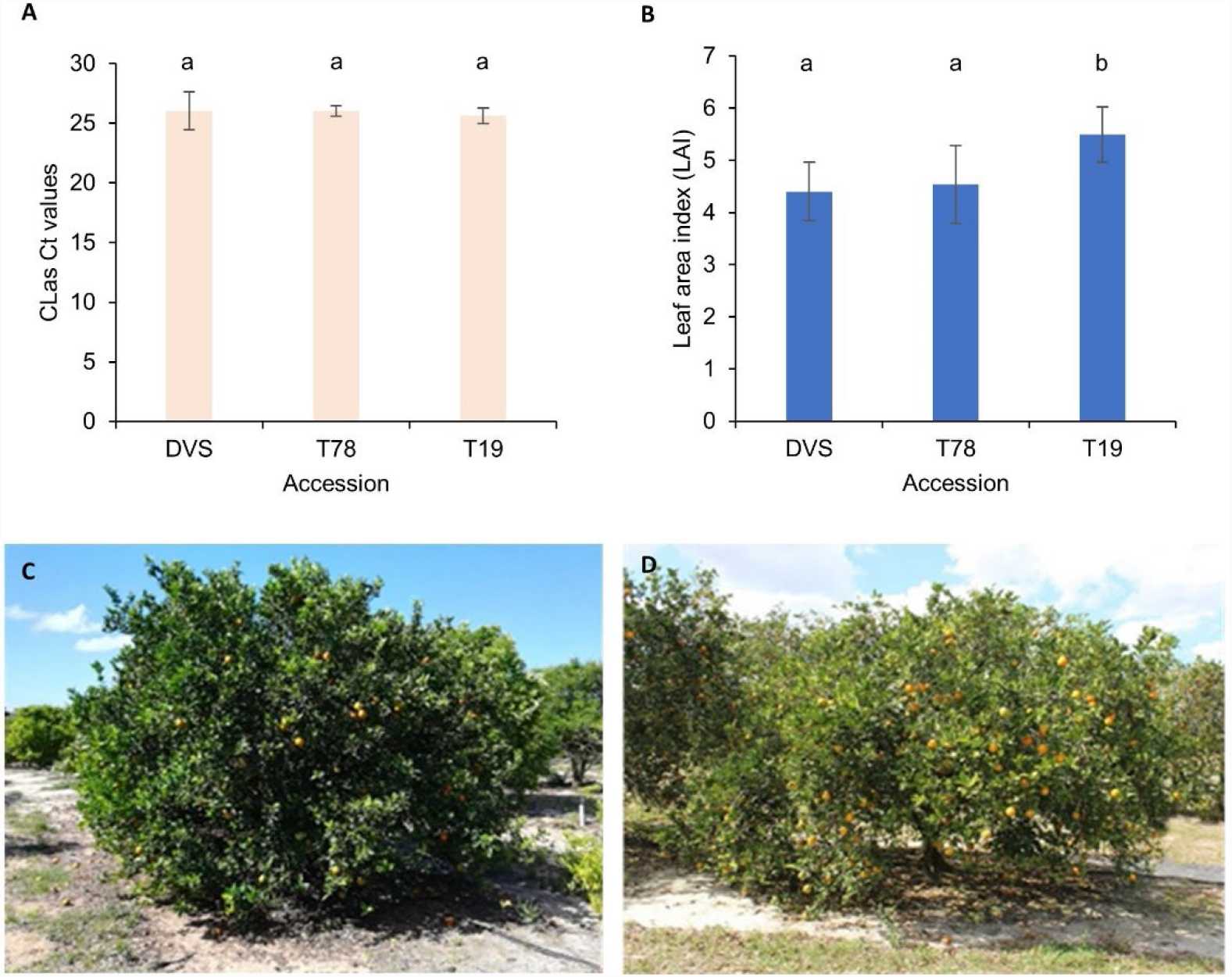
*C*Las titer and HLB symptoms on investigated Valencia sweet orange accessions. **A**. *C*Las titers of the DVS, T78, and T19 trees. The titer of *C*Las is represented by the CT values of qPCR tests using the 16S rRNA primers^66^. **B**. Leaf area indexes (LAI) of the investigated trees. The *C*Las titers and LAIs were tested on 5 to 6 different sections of each tree for 2 DVS, 2 T78, and 4 T19 trees. One largest and one smallest observed value of each tree were removed before calculating the mean CT and LAI values and their standard deviations (error bars in panels A and B). The lower-case letters a and b on top of the bars denote statistically different (p < 0.05) groups. **C and D**. Pictures of a T19 tree (C) and a DVS tree (D) taken in April 2019.

### Somatic structural mutations in the SWO accessions

We obtained no less than 280× PacBio CLRs for each of T19, SF, and T78. No lineage-specific mutation was detected in either T19 or SF, and three somatic SVs (TRA7, TRA44, and INS16) are shared by them (Supplementary Table 9), proving that SF was derived from the original T19 branch. TRA44 and TRA7 are complex TRAs both involving inversion and inter-chromosomal recombination (Extended Data Fig. 5). Four genes have been truncated by TRA7 and TRA44, and the promoter region of DVS7B01031 is translocated by TRA7 (Supplementary Table 10). INS16 is 5,086 bp Mutator–like transposable element (MULE) insertion on chr2A. Four lineage-specific mutations were identified in T78 (Supplementary Table 9 and Extended Data Fig. 5), including TRA22, a chromosomal recombination event between chr5B and chr1B; INV17, an 88,911 bp inversion on chr2B that truncated two genes; DEL58, a 1.8 Mb deletion on chr8B that has deleted 138 genes and truncated 1 gene; DUP52, an 5,617 bp tandem duplication on chr8A. TRA7, INV17, TRA22, TRA44, INS16, and DEL58 were verified through PCR amplification (Extended Data Fig. 6). We also detected forty SVs between DVS and both T19 and T78 that should have developed spontaneously (Supplementary Table 11 and Supplementary Note 2).

### Allele-specific expression and allelic expression ratio alteration in the SWO accessions

We configured an RNA-seq analysis pipeline to quantify the expression of genes both at the allelic levels (Extended Data Fig. 7) and carried out transcriptomic profiling for DVS, T19, and T78. In T19, 1,726 genes are significantly upregulated, and 1,503 downregulated compared with DVS and T78 (Supplementary Table 12). In T78, 1,054 upregulated and 1,161 downregulated genes are detected compared with DVS and T19 (Supplementary Table 13). For most genes with significantly altered expression in T19 (67.4%) or T78 (73.8%), the expression of their allelic genes is not significantly changed. Allele-specific expression (ASE) is observed in 32.4%, 34,0%, and 25.0% of the 10,737 tested DOGs in DVS, T19, and T78 transcriptomes (Fig. 3A). In 1,962 DOGs, ASE is detected in all three accessions (Supplementary Table 14). In most allelic expression ratio intervals, the proportion of genes is not significantly (p < 0.05 by Chi-square test) different among DVS, T19, and T78 (Fig. 3B), but 49.6% ± 0.07% allelic gene pairs have a ≥ 1.5-fold allelic expression difference in all the nine transcriptomes.

**Fig. 3:**
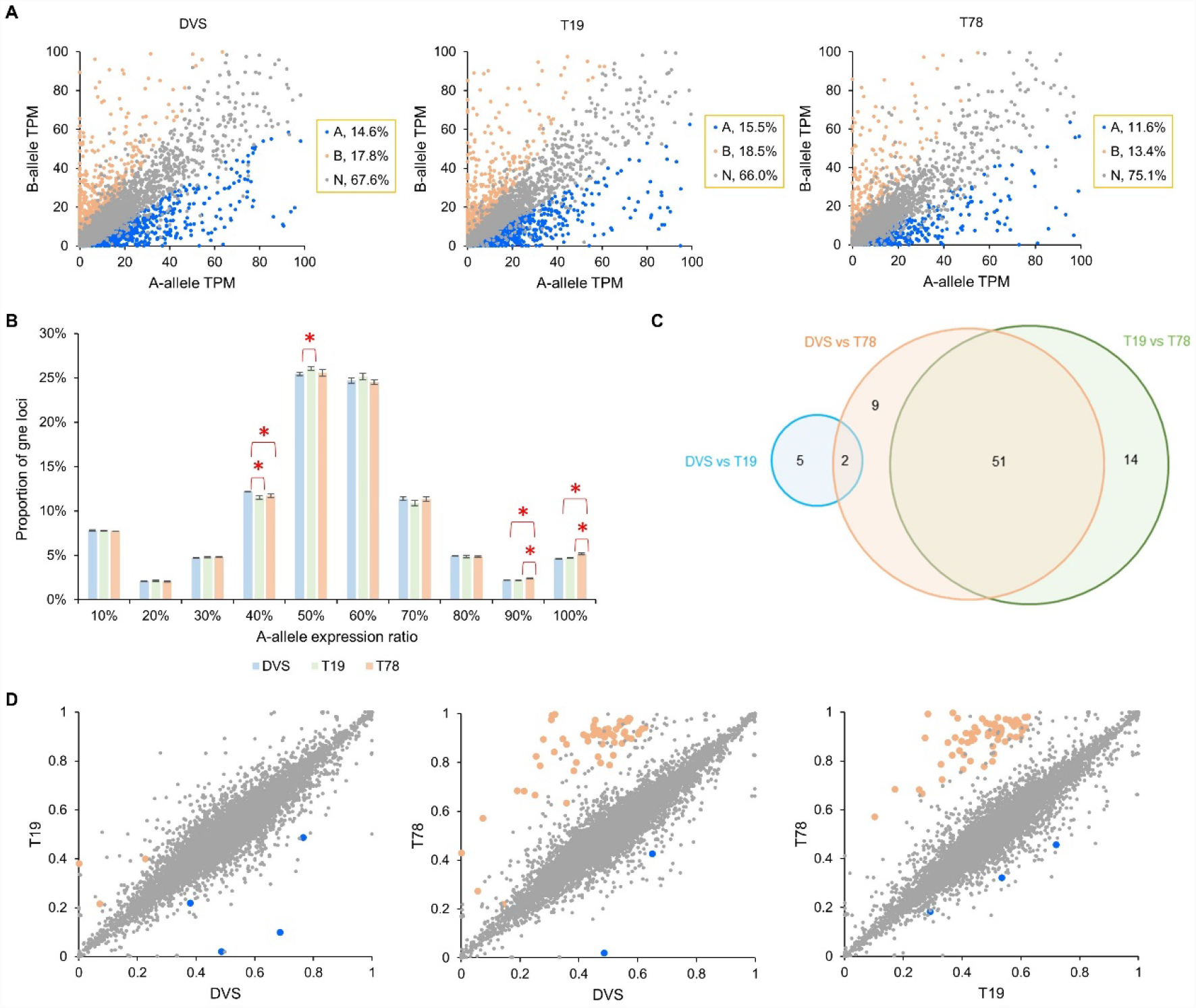
Allele-specific expression (ASE) and allelic expression ratio alteration (AERA) in DVS, T19, and T78. **A**. Allelic expression quantity distribution in DVS, T19, and T78 transcriptomes. Each dot represents a DVS orthologous group. A-allele and B-allele denote the corresponding allelic genes in DVS_A and DVS_B. The average normalized RNA-seq expression quantity TPM (transcripts per million) of the three transcriptomes was used for each sample. Genes with ASE biased towards A-allele and B-allele are highlighted with blue and light red colors, respectively. **B**. A-allele expression ratio distribution in the three sweet orange accessions. 10% denotes the closed interval [0%, 10%], 20% denotes the half-open interval (10%, 20%], and the rest are similar half-open intervals. Asterisks denote significant difference (^*^*p* < 0.05, two-tailed t-test) on the proportion of genes between the two samples pointed by the folding lines. **C**. The distribution of DVS orthologous groups (DOGs) with AERA among the three sweet orange accessions. **D**. Scatter plots showing the pairwise comparison of A-allele expression ratios among DVS, T19, and T78. The axes denote the A-allele expression ratio in the corresponding accessions, 0 indicates no A-allele expression, and 1 indicates only A-allele expression was detected. DOGs with a significant A-allele expression ratio increase and decrease in the horizontal axis accession compared to the vertical axis accession are colored blue and light red, respectively.

Allelic expression ratio alteration (AERA) is detected among DVS, T19, and T78 in 81 DOGs (Fig. 3C,D and Supplementary Table 15). Of the 76 DOGs with significant AERA in T78 compared to DVS and (or) T19, 62 contain genes located in the deleted 1.8 Mb segment (DEL58) of chr8B (Supplementary Table 15). The expression of the genes deleted by DEL58 is almost eliminated (Fig. 4A), while their orthologous genes are mostly (115/118) not significantly affected. The allelic expression ratio of DVS3A01852 interrupted by a 25,857 bp insertion (INS34 in Supplementary Table 11) in DVS has been reduced to 0.1%, compared to the 38.1% in T19 and 43.0% in T78 (Supplementary Table 15). On a few alleles directly affected by the somatic SVs, though no AERA is detected, their expression is significantly different in the mutant (Supplementary Table 10). In T19, the expression of DVS7B01006, truncated by TRA7, is significantly downregulated. The 3’ end of DVS3B03315 trimmed by TRA7 has significantly lower expression compared to DVS and T78, while its 5’ region is not significantly altered (Fig. 4B).

**Fig. 4:**
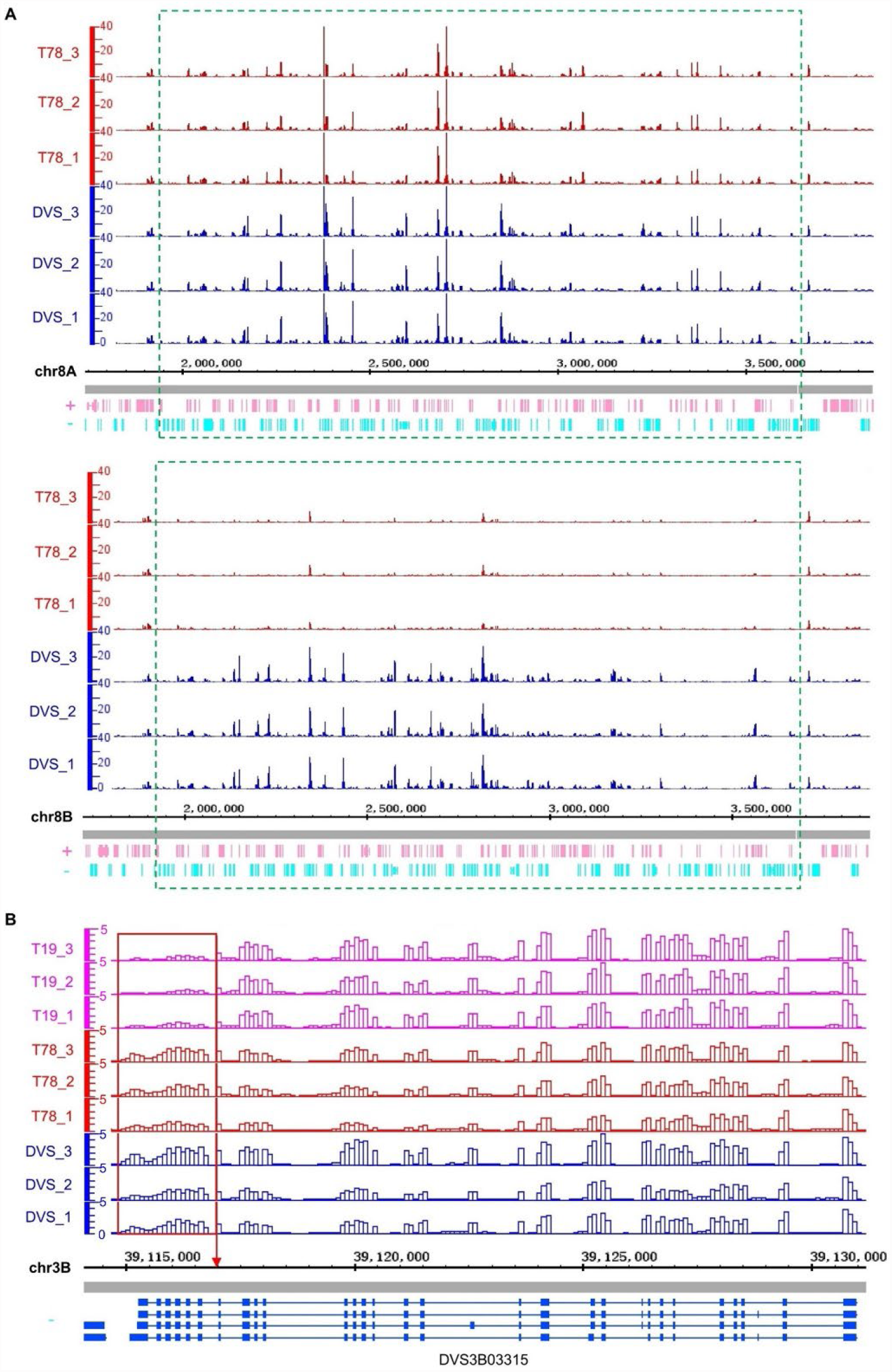
Allelic expression alteration related to structural mutations DEL58 and TRA7. The genomic region was split into 100 bp continuous non-overlapping windows. The vertical axes show the normalized uniquely mapped RNA-seq read counts in each window using the counts per million reads mapped (CPM) method. The horizontal axes denote the coordinates in the DVS genome. A. RNA-seq quantification in DEL58 (bottom panel) and its orthologous region (top panel) in DVS and T78. The green dashed-line frames denote the deleted region (bottom panel) and its orthologous region on chr8A (top panel). B. Regional expression alteration of DVS3B03315 truncated by TRA7. Different transcript isoforms are shown at the bottom, with blue rectangles representing exons and lines indicating introns. The red frame marks the truncated 3’ region of DVS3B03315 in T19, and the red arrow points to the chr3B:39,117,014 break endpoint of TRA7.

### Widespread heat shock protein upregulation in the HLB-tolerant mutant

The most significant alteration in the T19 transcriptomes is the strong upregulation of genes involved in protein folding (94 genes, FDR = 8.8E-37), response to heat (43 genes, FDR = 8.5E-26), and ribosome biogenesis (65 genes, FDR = 1.9E-11) (Supplementary Table 16). The most upregulated genes involved in protein folding or response to heat were *HSPs*. Further analysis reveals that more than half of HSP genes (68/133, enriched by19.6 fold and P = 7.0E-51) are significantly upregulated in T19 by 2^2.6 ± 1.1^-fold compared to DVS and T78 (Fig. 5A), including multiple *HSP60*s, *HSP70*s, *HSP90*s, and small *HSP*s (Supplementary Table 17).

**Fig. 5:**
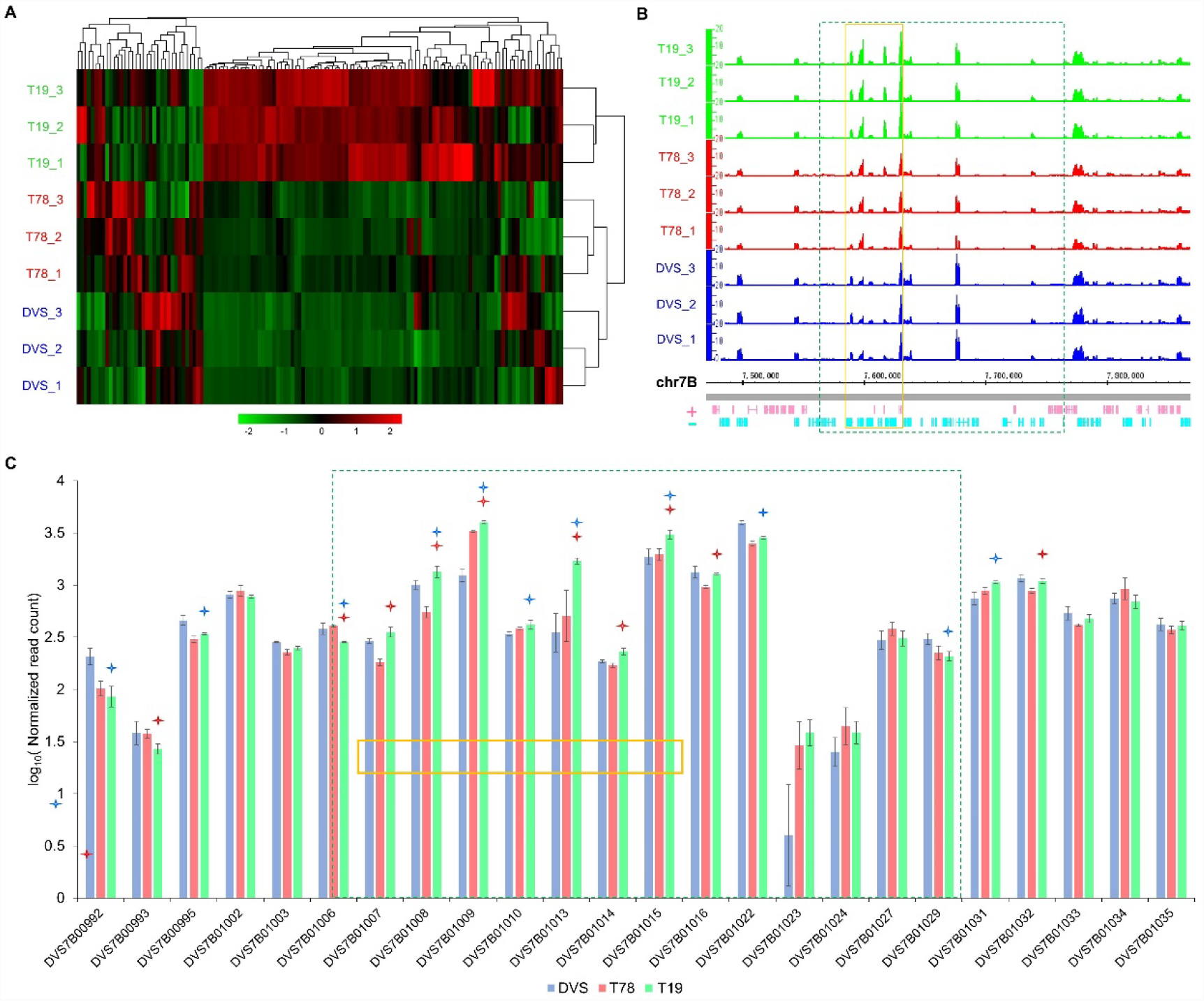
Putative transcriptome alteration related to the high HLB tolerance of T19. A. Heat map of RNA-seq expression quantification of heat shock protein (HSP) encoding genes. Each column shows the normalized expression quantity of an HSP encoding gene in the nine transcriptomes shown on the left. The dendrograms depict the results of hierarchical clustering based on Euclidean distance among the transcriptomes (right) and the HSP genes (top). B. RNA-seq read abundance surrounding the translocated region of TRA7. The plot was drawn using the same method for Fig. 4, with the vertical axis denoting the read abundance in CPM and the horizontal axis denoting the coordinates in the DVS genome. C. Histogram of the normalized gene expression around TRA7. Only genes with a minimum average read count of 3 reads per transcriptome have been shown. On top of the green bars, blue and red stars denote being significantly (FDR < 0.1) different from DVS and T78 in T19, respectively. In B and C, the green dashed-frames denote the translocated chr7B:7,571,519-7,765,552 region of TRA7, and the orange frames highlight the seven genes with upregulated expression (*p* < 0.05) compared with both DVS and T78.

Upstream and downstream regions of SV breakpoints were scanned for those enriched in genes with significantly altered expression. The results show that the upregulated genes are significantly overrepresented (FDR < 0.05) at one terminal of the segment (blue arrow of TRA7 in Extended Data Fig. 5) translocated by TRA7 in T19 (Fig. 5B). Seven genes within that region are upregulated compared to DVS and T78 (Fig. 5C). Three of the seven genes are putatively related to the upregulation of HSP encoding genes, which are *CsDNAJ* (DVS7B01009)^16^, *CsHSP17*.*4B* (DVS7B01013)^17^, and *CsCEBPZ* (DVS7B01014)^18,19^.

## Discussion

We assembled a phased chromosome-level sweet orange reference genome with improved completeness, accuracy, and gene structure annotation compared to the recently updated HSO genome^4^. The completeness and RNA-seq data mapping rate test results (Extended Data Fig. 8 and Supplementary Note 3) suggest that DVS should be the reference of choice for SWO, cultivated mandarins, grapefruits, and other inter-specific hybrids between mandarin and pummelo. This phased assembly also has brought an end to the long-disputed genetic composition and parental hybridization processes of SWO (Supplementary Note 4). The intra-genomic variants detected in DVS outnumbered the previous studies using mapping-based methods with NGS data^2,3^. We also observed a vast number of monoallelic genes in the DVS genome, indicating its genetic redundancy has been discounted considerably. Previously, ASE was only detected on individual SNVs in SWO^20^. With DVS as the reference, the gene expression of SWO could be quantified directly at the allelic level using RNA-seq data. DVS could also be applied in other allelic-level studies, including allele-specific protein expression (ASPE) analysis^21^ and allele-specific genetic engineering^22^. Many citrus cultivars contain genetic materials from both pummelo and mandarin^3,5,23^, making DVS very important in the genus.

As the length of sequencing reads improved^24^, the high intra-genomic heterozygosity, such as found in SWO, could be utilized as an advantage in phased genome assembly. We adapted the assembly parameters to the allelic difference level of SWO and assembled most genomic regions directly into two haploid contigs. Our method avoided mapping-based phasing in the highly heterozygous genomic regions and has achieved a very low hamming error rate^25,26^. Genetic maps^2,3,27^ or the Hi-C technology^4,28^ have been utilized in the scaffolding step of the citrus genome assembly. This study shows that for genomes with diverged repeat units among long-tandem repeat regions, the long sequencing reads could have contained enough information for chromosome-level scaffolding. CANU’s capacity to resolve high-similarity repeats^29^ is important for the successful application of our method.

Most irradiation-induced and spontaneous somatic SVs have developed through different molecular mechanisms. Irradiation could cause a wide variety of DNA lesions, among which double-strand breaks are the most relevant to structural mutations^30,31^. In this study, five of the seven putative irradiation-induced SVs in T19 and T78 involved double-strand break repair in the non-homologous end-joining manner^32^. TE activity has been reported to be the primary source of spontaneous SVs in a few plants^33,34^. A few bud sports from sweet orange and other citrus species did have been distinguished by TE polymorphism^35,36^. In 114 SWO accessions, TE insertions accounted for 40.1% of the large somatic insertions detected^4^. This study shows that the spontaneous SVs detected are mainly (29/40) derived from the insertion of three MULEs (Supplementary Table 11), indicating they might be hyperactive and have played an essential role in the formation of some SWO cultivars.

SVs have been linked with gene expression differences in a few species^37–39^. The retrotransposon insertion in the promoter region caused the *Ruby* gene to be expressed under cold stress in blood SWO^7^. Diversified effects of SVs on gene expression have been observed in this study. The expression of deleted allelic genes in T78 was generally eliminated as expected, and their orthologous genes’ expression was mostly unaffected. The case for a truncated gene was more complicated, which could either be entirely downregulated or only have a 3’ end downregulation. Most chromosomal rearrangements in T19 and T78 had no noticeable impact on the expression of adjacent genes except for TRA7. Chromosomal rearrangements could affect the expression of adjacent genes by causing changes in chromatin topology, but their effects remain difficult to predict^40^.

Dysfunctional phloem induced by *C*Las infection is one of the primary causes of HLB symptoms. Multiple mechanisms have been related to the enhanced HLB tolerance/resistance (Extended Data Fig. 9), including lowered *C*Las titer through enhanced NPR1-dependent defense^41,42^ or overexpressed antimicrobial peptides^43–45^, enhanced phloem cellular homeostasis^46–48^, and more vigorous phloem regeneration^15,49^. Through summarizing previous studies, the HLB symptom development is proposed to include six steps, and we found the proposed HLB tolerance/resistance mechanisms could be associated with the symptom development steps in this disease (Extended Data Fig. 9). The long-lasting HLB tolerance of T19 should be mainly ascribed to the upregulation of multiple classes of *HSP*s (the mechanism B in Extended Data Fig. 9), which promotes phloem homeostasis and inhibits subsequent HLB symptom development. HSPs primarily function as molecular chaperones and are the key components responsible for keeping cellular protein homeostasis under abiotic and biotic stresses^50,51^. HSPs play important roles in inhibiting programmed cell death in animals^52,53^ and plants^54–57^. Multiple classes of HSPs have been reported to counter reactive oxygen species (ROS)^58,59^, a key regulator of plant programmed cell death^60^. Downregulation of a few HSPs has been reported in HLB susceptible citrus after *C*Las infection^46,61–63^, while upregulation of them was putatively related to enhanced HLB tolerance^46^. Ma et al. (2022) inferred the accumulation of reactive oxygen species (ROS) to be caused by chronic immune responses in the phloem tissue^64^. However, there is no proof that *C*Las could be recognized by citrus within the phloem, which is a prerequisite for inducing immune responses. Thus, we hypothesize that the *C*Las propagation breaks the cellular homeostasis, causing protein aggregation, ROS accumulation, and sequentially programmed cell death in the phloem, then the released *C*Las cells induce the pathogen/microbe-associated molecular pattern-triggered immunity responses^65^.

## Methods

### Plant materials

Valencia sweet orange buds were exposed to 50 Gy gamma irradiation, and more than 1200 trees were produced by budding onto Volkamer lemon rootstock seedlings and planted in the field in 1992, as part of a mutation breeding project. From these trees, 6 were identified as bearing nearly seedless fruit including T19, SF, and T78. These were subsequently repropagated onto Carrizo citrange rootstock and at least 2 of T78 and 3 of T19 were planted in the field near Lake Alfred, FL in summer 2000. They along with DVS have been grown in the field under the same management conditions since then DVS and OVS are different Valencia orange trees, the former being the tree used to produce the genome assembly reported here, and the latter being the budwood source for the production of the irradiated Valencia population from which T19, T78, and SF were selected. The SF tree was known to be grafted from one of the original nearly seedless selections from the same experiment, but the identification tag was lost, so its clonal identity was uncertain. HLB was first detected in this field location in 2008, and by 2010 virtually all trees were showing symptoms of infection. These specific individuals, although also exhibiting HLB symptoms, were first noted for their obvious superior performance and substantially better appearance compared with all other nearby trees of standard Valencia that were in severe decline, as well as a wide range of other materials from the breeding program likewise in severe decline, in 2017. T19 trees have retained their tolerant phenotype, but T78 trees have gradually declined since first noted.

### Measurement of leaf area index and *C*Las titer

The leaf area index (LAI) was measured using AccuPAR LP-80 (Meter Group, Pullman, WA, USA) near solar noon in June 2021. The external photosynthetic active irradiation (PAR) sensor was placed in a nearby open area, and the LP-80 instrument PAR probe was placed under the canopy of each tested tree. The LP-80 computed LAI from the PAR readings and χ (leaf angle distribution parameter). The default χ parameter (χ = 1) was applied. On average, 6-7 measurements per tree were taken around each tree.

For *C*Las titer measurement, DNA was extracted from leaf midribs and petioles of each tree using the Plant DNeasy Mini Kit (Qiagen, Valencia, CA, USA) according to the manufacturer’s instructions. qPCR quantification of the *C*Las titer using 16S rRNA primers was carried out as described by Li et al. (2006)^66^. qPCR was performed on an Agilent Mx3005P (Agilent Technology Inc, Waldbronn, Germany) real-time PCR system with the Brilliant III Ultra-Fast QPCR Master Mix (Agilent Technology Inc, Waldbronn, Germany).

### DNA and RNA extraction

For next-generation sequencing (NGS) and PacBio sequencing, young leaves of DVS, T78, T19, and SF (PacBio sequencing only) were collected from new flushes in April 2018 and April 2019, respectively. We used the CTAB method^67^ to extract genomic DNA for NGS. For PacBio sequencing, genomic DNA was isolated using Nanobind Plant Nuclei Big DNA Kit (Circulomics Inc., Baltimore, MD, USA) following its manufacturer’s instructions.

As for RNA extraction, mature leaves were collected from three different tree parts for DVS, T19, and T78 as replicates. In total, 9 RNA samples were extracted using TRIzol(tm) and RNA Purification Kit following the manufacturer’s protocol (ThermoFisher, Waltham, MA, USA). RNA was further purified using the TURBO DNA-free(tm) kit (ThermoFisher, Waltham, MA, USA) to eliminate genomic DNA. Both NanoDrop Spectrophotometer (NanoDrop Technologies, Wilmington, DE, USA) and Agilent 2100 Bioanalyzer (Agilent Technologies, Waldbronn, Germany) were used to assess the RNA quality and quantity.

### De novo assembly of DVS and the mutants

Whole-genome PacBio continuous long reads (CLR) were obtained for DVS, SF, T78, and T19 on the PacBio Sequel II system (Pacific Biosciences, Menlo Park, USA). One hundred-fold coverage of corrected reads (N50=42.0 kb) was used. A minimum of 98.5% overlap identity was required in the assembly step to reduce collapsed assembly in heterozygous regions. We carried out de novo assembly of DVS using MECAT2^68^ with four different minimum read overlap lengths (500 bp, 2 kb, 5 kb, and 10 kb). A 2 kb minimum read overlap length was identified as optimal since the assembly achieved the second-largest N50 and the largest assembly size. The assemblies’ accumulative length and contig N50 were assessed using QUAST v5.1^69^. De novo assembly of the three mutants was carried out by MECAT2 using the optimal parameters observed for DVS (Supplementary Table 18).

We also carried out de novo assembly of DVS using CANU v2.1^29^ with the same optimal settings. CANU produced ∼26.6 Mb more sequences mainly derived from repetitive regions. The contigs in the CANU assembly were connected into pseudochromosomes through (1) phased assembly of the collapsed and expanded regions by FALCON and FALCON-UNZIP in pb-falcon v2.24^70^; (2) resolving the repeat units in long tandem repeat regions (Extended Data Fig. 1). In the process, each contig in the CANU assembly was mapped against the remaining contigs using minimap2 v2.17^71^ to detect orthologous regions. The sequencing depth across the assembly was output using BEDTools v2.29.2^72^ in 1 kb windows. The unphased regions and putatively collapsed regions were subjected to phased assembly. A circular mitochondrion genome and a circular plastid genome were manually recovered by aligning and connecting several contigs with high coverage. The chromosomes were named in concordance with the haploid CCL genome^3^. Using the CLR reads, three rounds of polishing were carried out by pbmm2 and arrow in GenomicConsensus v2.3.3 (Pacific Biosciences, Menlo Park, USA).

### Hamming error rate estimation

The paternal and maternal parents of SWO are unknown, so we estimated the hamming error rate across the genome based on its di-haploid offspring HSO^4^; 100 × sequencing reads were simulated without sequencing error for the HSO genome by wgsim (acquired on 06/15/2021 from https://github.com/lh3/wgsim). The simulated reads were mapped to DVS using minimap2 v2.17. The simulated reads uniquely mapped to DVS_A (*C*_*A*_) and DVS_B (*C*_*B*_) were counted in 20 kb continuous windows with 15 kb overlap across the HSO genome. The hamming error rate was calculated as the minimum of *C*_*A*_ and *C*_*B*_ divided by the sum of *C*_*A*_ and *C*_*B*_ in each window. Because HSO is a di-haploid offspring of SWO, the windows overlapping with putative chromosomal recombination loci in HSO, identified with surrounding windows switching from *C*_*A*_ *> C*_B_ to *C*_*B*_ *> C*_*A*_ or vice versa, were excluded from the calculation.

### Genetic origin inference of the DVS chromosomes and switch error detection in interspecific regions

Twenty mandarin and twenty pummelo whole-genome NGS data sets (Supplementary Table 3) were downloaded from the NCBI database. The DVS assembly was randomly separated into two sets, each including chromosomes 1 to 9. The two chromosome sets were analyzed separately. First, the mandarin and pummelo NGS data were mapped to a chromosome set by BWA v0.7.17^73^. Small variants and genotypes of the 40 samples were called using SAMtools v1.10^74^ and BCFtools v1.10^75^. The genotypes were filtered by requiring a minimum genotype quality of 30 and local sequencing depth not exceeding 1.5-fold of the average sequencing depth of the sample. The reference allele was denoted ‘0’, and ‘1’ denoted the alternative allele in the genotypes. The variants with only homozygous genotypes or those with < 10 samples genotyped in either species were excluded.

The distances between sample *i* and the reference were calculated in continuous 10 kb windows as:

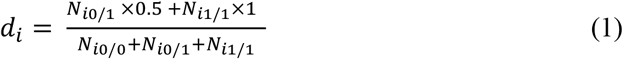

The symbols in the formula: *d*_*i*_, the distance between sample *i* and the chromosome set in the analyzed window; *N*_*i0/1*_, *N*_*i0/0*_, and *N*_*i1/1*_ denote the number of variants with 0/1, 0/0, and 1/1 genotypes in sample *i*. A minimum of 10 variants was required to be genotyped for a sample to be counted in a window. One largest and one smallest distance were removed from either species in each window. When ≥ 10 samples remained available in both species, a t-test was carried out, and the reference window would be assigned to the species with significantly (*p* < 0.05 by two-tailed t-test) smaller distances. For windows with not enough variants genotyped in one group, the read coverages (normalized by whole-genome sequencing depth) were compared between the two species, and the reference window would be assigned to the species with significantly (*p* < 0.05 by two-tailed t-test) higher coverage. The genetic origin of repetitive regions and regions with sparse variants was inferred based on their adjacent windows. A switch error was detected when P-M and M-P switches were observed simultaneously on DVS_A and DVS_B in the corresponding orthologous windows.

### Quality evaluation and comparison between DVS and HSO

HSO and DVS were aligned in a pairwise fashion via minimap2 v2.17. The unaligned regions were output using BEDTools v2.29.2. The whole-genome dot-plot was drawn by D-GENIES v1.2.0^76^ using the whole-genome alignment with HSO as the reference. For assembly quality assessment, we obtained 26–30 × pair-end sequencing reads (150 bp × 2) from Illumina HiSeq 2500 (Illumina, San Diego, CA, USA) for DVS, T19, and T78. The K-mer based completeness and the error rates of the assemblies were assessed by Merqury v1.3^77^. Core gene completeness test was carried out by BUSCO v5.0.0^78^ using its eudicots_odb10.2019-11-20 database.

To test the mapping rate difference with DVS and HSO, both with organelle genomes included, we downloaded twelve whole-genome NGS data sets (Supplementary Table 2) of different sweet orange cultivars from NCBI. Then we aligned them to DVS and HSO by BWA v0.7.17 using the default parameters. The mapping rates and properly paired mapping rates were output using SAMtools v1.10.

To analyze the sequencing depth homogeneity across the HSO assembly, we downloaded its whole-genome CLR reads (SRR5838837) from NCBI^4^. The CLR reads of DVS and HSO were mapped to their respective assemblies by minimap2 v2.17. We output the whole-genome sequencing depths across the two assemblies using BEDTools v.2.29.2, which were normalized by being divided by their whole-genome average sequencing depth.

### Intra-genomic variation detection and annotation

We aligned DVS_B to DVS_A using minimap2 v2.17 and called the small variants using BCFtools v1.10 with the consensus model^75^. Only regions with one-to-one unique alignments were used to call the variations. SNVs or indels within 10 bp distance of another indel were filtered. For SV detection, we carried out whole-genome alignment on DVS_A and DVS_B using Mummer v4.0.0^79^, and the SVs between DVS_A and DVS_B were called by MUM&CO v2.4.2^80^. The circular graph (Fig. 1A) showing the distribution of the variations was drawn using Circos v0.69-9^81^. Genetic variant annotation and functional effect prediction for the small variants were predicted by SnpEff v5.0^82^, which defined the categories of high, moderate, low, and modifier impact variants. BEDTools v2.29.2 was used to detect genes overlapped with all the SV types.

### Annotation of the DVS genome

Putative TEs were predicted in the DVS genome using both RepeatModeler v2.0.1^83^ / RepeatMasker v4.1.1^84^ and the Extensive de-novo TE Annotator (EDTA) v1.8.3 pipeline^85^. Unclassified TEs were subjected to convolutional neural networks (CNNs) based classification by DeepTE (acquired from https://github.com/LiLabAtVT/DeepTE on 01/15/2021) using the plant model^86^.

The DVS genome was soft masked using RepeatMasker v4.1.1 based on the non-redundant TE library from the EDTA pipeline. Ab initio gene prediction and transcriptome data assembly-based methods were applied in combination to annotate gene models in the genome. Eighty SWO transcriptomic RNA-seq data (Supplementary Table 19) from multiple tissue types were downloaded from NCBI and mapped to DVS using HISAT2 v2.2.1^87^ for RNA-seq evidence in ab initio annotation. The UniProtKB/Swiss-Prot plant database (accessed on 10/13/2020)^88^ was used to generate protein hints with GenomeThreader v1.7.3^89^. Then GeneMark-EP+ v4.65^90^ and Augustus v3.4.0^91^ were trained based on the RNA-seq and protein hints using the BRAKER v2.1.5 pipeline^92^. For transcriptome assembly-based annotation, the RNA-seq reads uniquely mapped to DVS_A or DVS_B were separated into two datasets. The reads with equal mapping scores to DVS_A and DVS_B were added to both of them. RNA-seq read assembly and protein-encoding transcript selection were then carried out separately for DVS_A and DVS_B with the Mikado v2.0 pipeline^93^.

### Gene annotation and RNA-seq mapping rate comparisons among citrus assemblies

To compare the completeness of gene structure annotations, the protein sequences encoded by the first transcript of all genes were output for DVS, DVS_A, DVS_B, HSO^4^, CCL^3^, HKC (*Atlantia buxifolia*), XJC (*Citrus ichangensis*), XZ (*Citrus medica*), HWB^94^, MSYJ^6^, HK (*Citrus japonica*)^95^, ZK (*Poncirus trifoliata*)^27^, and PTR^28^. The protein sequences were tested against the eudicot core gene set (eudicots_odb10.2019-11-20) by BUSCO v5.0.0 using the protein mode^78^.

The RNA-seq data mapping rate was compared with different citrus assemblies as the reference. A masked DVS version was generated by masking the allelic genes with < 3 SNVs / kb in the exonic regions except for one allele in DVS_A. The separately tested DVS_A and DVS_B were not masked. Forty RNA-seq data sets from SWO, grapefruit, mandarin and pummelo were downloaded from NCBI (Supplementary Table 20). The RNA-seq data were mapped to the assemblies by HISAT2 v2.2.1, which reported the overall, concordant, and unique mapping rates.

### Analysis of orthologous gene groups in citrus

Proteins encoded by the first transcript of all gene models in DVS_A, DVS_B, HSO, CCL, HWB, MSYJ, HK, and PTR were output with GffRead v0.12.1^96^. The protein sequences were clustered into citrus orthogroups using OrthoFinder v2.5.2, which applied a phylogeny-based method^97^. The first protein in each COG was searched against the PANTHER v16^98^ database using MMseqs2 v12-113e3^99^. HQ-COGs were identified by requiring two conditions: (1) including members from at least three citrus genomes; (2) having at least one target hit in the PANTHER database with the alignment covering ≥ 30% of both the query and the target sequences and an E-value < 1.0E-3; those not meeting the criteria were classified as LQ-COGs. Co-synteny orthologous genes between DVS_A and DVS_B were inferred using MCScanX^100^.

### Detection of somatic SVs in the radiation-induced mutants

Assembly and mapping-based strategies were integrated to detect somatic structural variants in SF, T19, and T78. The assemblies of the mutants were aligned to the DVS assembly, and candidate SVs were called by MUMMER v4.0.0. The four MECAT assemblies of DVS were also aligned to the DVS genome using the same method to acquire false-positive SVs. In the mapping-based method, the CLRs were mapped to DVS with Minimap v2.17, and SV calling was carried out by Sniffles v1.0.12^101^. A filter requiring a minimum of 10 zero-mode waveguides support was applied on the mapping-based SVs. The results of the two strategies were compared, and the common ones were selected for further analysis. Maximum margin distances of 50 bp for breakpoint ends of TRAs, and 50 bp or 10% of the SV lengths (whichever smaller) for DELs, INSs, DUPs, and INVs, were allowed for the SVs from the two strategies to be considered the same. We designed primers and carried out PCR verification of 6 SVs (Supplementary Table 21).

### RNA-seq and differential expression analysis

Whole-transcriptome sequencing, including rRNA depletion and stranded library construction, was carried out by BGI Genomics (Shenzhen, China). The sequencing via Illumina HiSeq 4000 (Illumina, San Diego, CA, USA) produced > 5 Gb clean pair-end (150 × 2) reads for each sample. The masked DVS in RNA-seq mapping rate tests was used as the reference in RNA-seq analysis. Salmon v1.4.0^102^ was applied in read mapping and counting, with multi-mapped reads assigned by the expectation-maximization algorithm^103^. Read count normalization and differential expression tests were carried out by DESeq2 v1.30.1^104^. Genes with significantly (FDR < 0.1) differential expression at the allelic level were first detected in a pairwise manner among DVS, T19, and T78. Then those significantly upregulated or downregulated genes compared to both DVS and T78 (T19) were identified as differentially expressed genes in T19 (T78).

To visualize the abundance of RNA-seq reads in the SV-affected regions, we mapped all RNA-seq data to DVS using HISAT2 v2.2.1. The uniquely mapped reads were counted in 100 bp continuous non-overlapping windows across the DVS genome using deepTools v3.5.0^105^ with the CPM normalization method. The read abundance in regions of interest was visualized using the Integrative Genomics Viewer (IGV) v2.8.0^106^.

### Analysis of allele-specific expression and allele-expression ratio alteration

Based on the RNA-seq data of DVS, T19, and T78, the TPM (transcripts per million) normalized expression quantity of allelic genes were calculated by Salmon v1.4.0 and tximport v1.16.0^107^. Low expression orthologous groups with < 0.25 transcripts per transcriptome were filtered. For each genotype, allele-specific expression (ASE) was detected on a gene locus if the two alleles had mean TPM values differing by ≥ 1.5 fold in the three transcriptomes, and the FDR is smaller than 0.05 by two-tailed t-test. To detect allelic expression ratio alteration (AERA) among the accessions, we compared the A-allele (allelic genes in DVS_A) expression ratios, calculated as [A-allele TPM / (A-allele TPM + B-allele TPM)], on each gene locus among DVS, T19, and T78. If the mean A-allele ratio was different by ≥ 1.5 fold and the FDR was < 0.05 by two-tailed t-test, AERA was inferred on the gene locus between the two accessions.

### qPCR quantification of gene expression

Twenty-three DEGs in T19 (Supplementary Tables 22 and 23) identified by RNA-seq were selected for quantitative real-time PCR (qPCR) verification. First-strand cDNA was synthesized from 0.3 μg of total RNA using the Affinityscript qPCR cDNA Synthesis Kit (Agilent Technologies, Santa Clara, CA, US). qPCR was performed using the Brilliant III Ultra-Fast SYBR Green QPCR Master Mix (Agilent Technologies, Santa Clara, US) following its instructions. With 18S rRNA as the reference gene^108^, the 2^-ΔΔCt^ Ct method was applied to analyze the qRT-PCR results^109^.

## Data availability

All sequencing data generated in this study (PacBio sequencing, NGS sequencing, and RNA-seq data) have been deposited in the National Center for Biotechnology Information (NCBI) under BioProject ID PRJNA735893. The genome assembly and gene annotation of DVS have been submitted to NCBI under the genome Accession IDs JAHMIS000000000 (DVS_A) and JAHMIT000000000 (DVS_B). Data supporting the findings of this work are available within the paper and its Supplementary Information files.

## Acknowledgements

This work was supported in part by the U.S. National Institute of Food and Agriculture (NIFA) under Grant 2017-70016-26051 to F. L. and F. G. and the U.S. National Science Foundation (NSF) under Grant ABI-1759856 and MTM2-2025541 to F.L. This research was partially supported by grants from the Citrus Research and Development Foundation (CRDF 15-010, CRDF RMC 18-010, and CRDF RMC 18-011), and the New Varieties Development and Management Corporation to F.G.

## Author contributions

F.G. and F.L. conceived and designed this project. F.G. and Q.B.Y. observed and assessed the HLB tolerance of the plants. Q.B.Y carried out all the biological experiments. B.W. did all the bioinformatical and statistical analyses. B.W., Q.B.Y., F.G., and F.L. wrote the paper. Z.N.D and Y.P.D evaluated the plant materials and the data analysis results. All authors have read and approved the final version of this paper.

## Figure Legends

**Extended Data Fig. 1:**
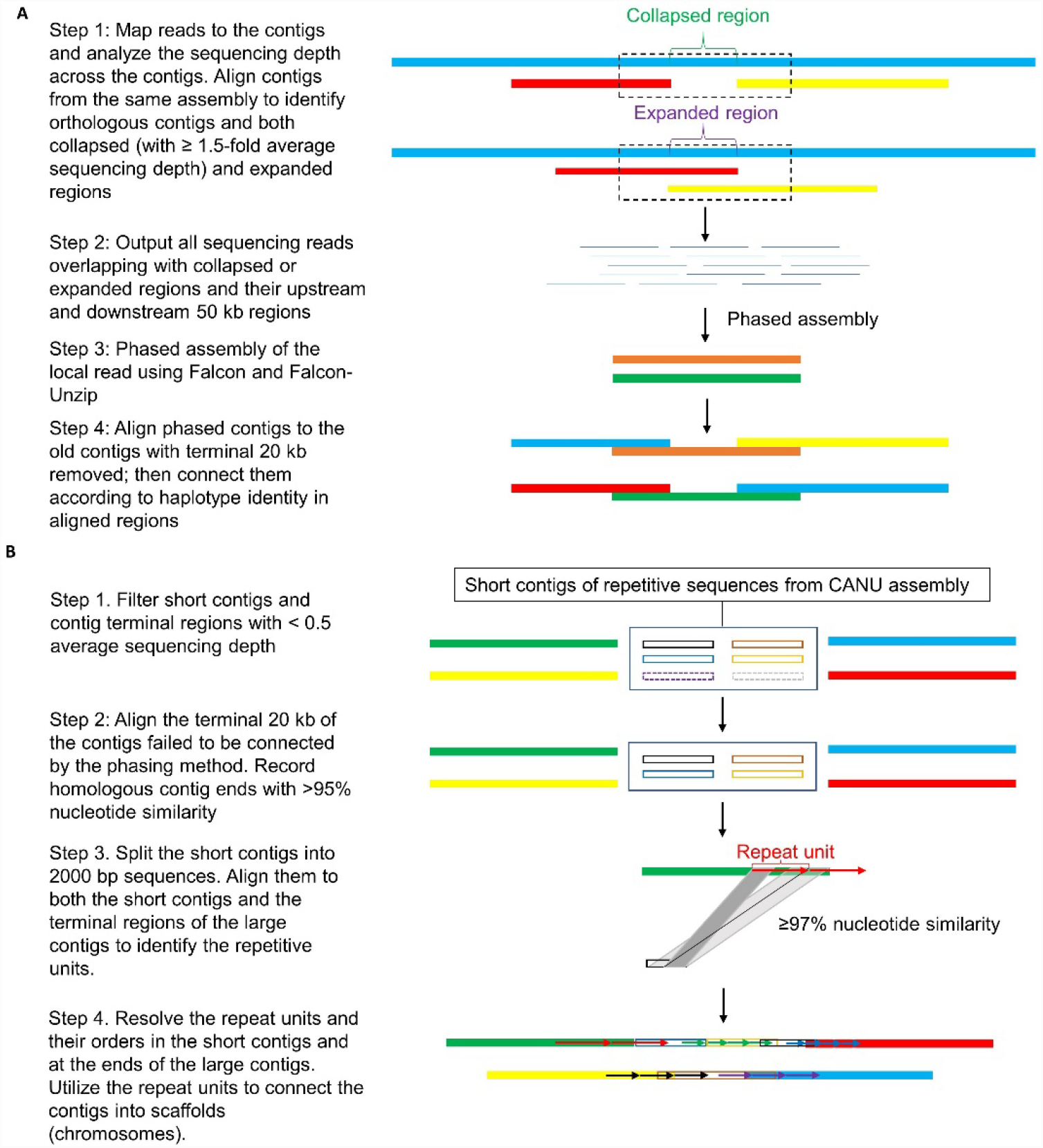
Scaffolding the contigs through phasing and repeat unit resolving in DVS assembly. **A**. Connecting contigs and fixing putative switch errors through the phased assembly of the collapsed and expanded regions. **B**. Scaffolding the contigs through resolving the repeat units. CANU could separate repeats with > 3% difference^59^. Thus ≥ 97% nucleotide similarity was required to identify regions composed of the same repeat units. The arrows with different colors indicate different repeat units.

**Extended Data Fig. 2:**
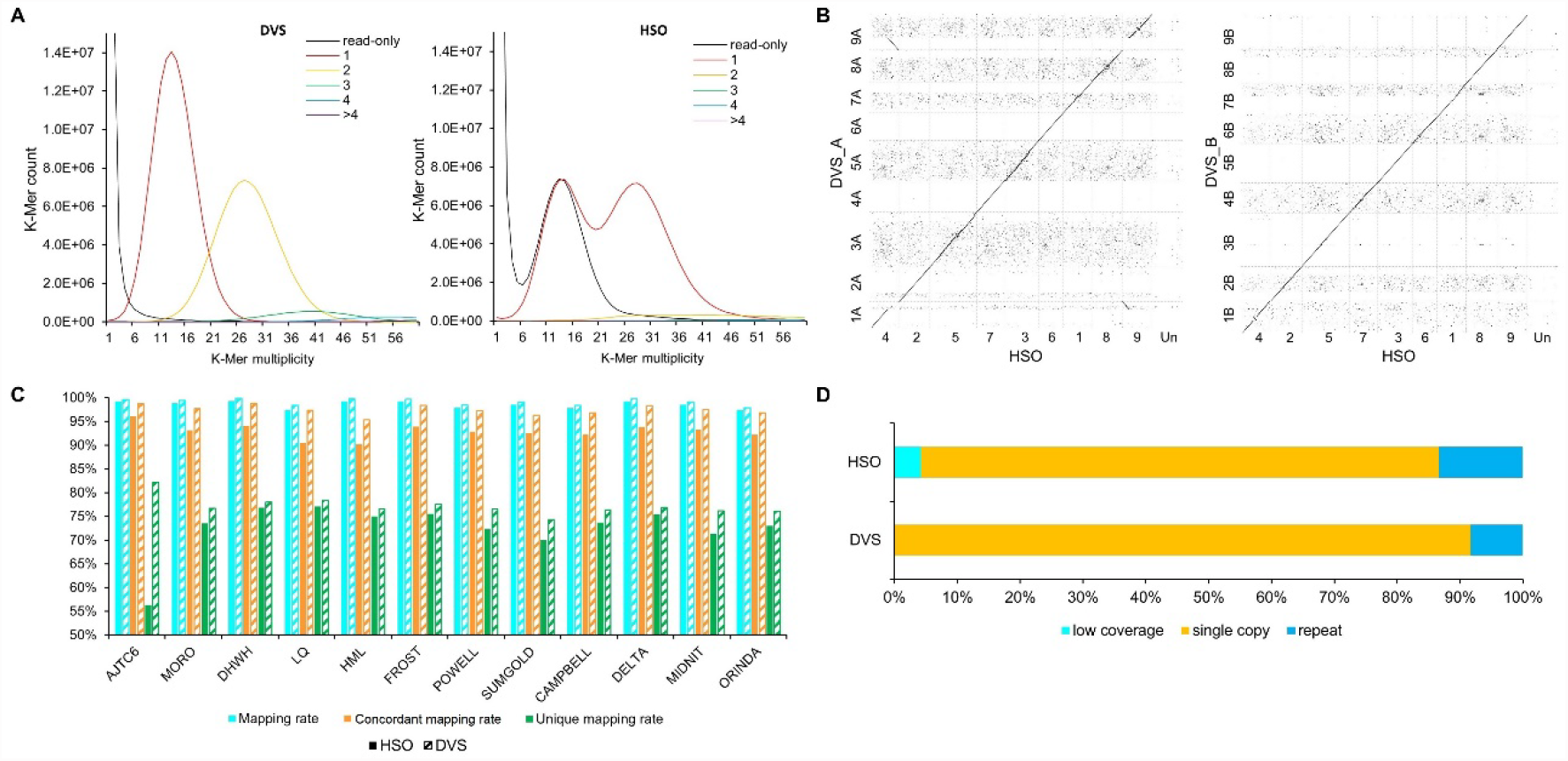
Comparison between HSO and DVS assemblies. **A**. Distribution of K-mer counts in SWO whole-genome NGS data per copy numbers found in the assemblies. The colors of the curves indicate the copy number of the K-mer in the assembly as shown in the legend. ‘read-only’ indicates the K-mer was only detected in the NGS reads. **B**. Whole-genome dot-plot comparison between HSO and chromosomal sets DVS_A and DVS_B. Regions of homology are plotted as diagonal lines or dots. The D-GENIES program split the query assemblies (DVS_A and DVS_B) into ten mega-base chunks before alignment^76^. Regions aligned in large continuous fragments have fewer noisy dots. **C**. Whole-genome NGS data mapping rates with HSO and DVS as the reference. Pair-end sequencing reads of 12 sweet oranges (Supplementary Table 2) were mapped to HSO and DVS with the same alignment parameters. The mapping rate is the proportion of individual reads mapped regardless of the mapping status of their paired reads. The concordant mapping rate is the ratio of reads mapped in proper pairs. Unique mapping rate is the ratio of uniquely mapped reads. **D**. Proportions of low coverage (≤ 0.5 fold of sequencing depth), single copy (> 0.5 fold and < 1.5 fold), and repeat regions (≥ 1.5 fold) in HSO and DVS.

**Extended Data Fig. 3:**
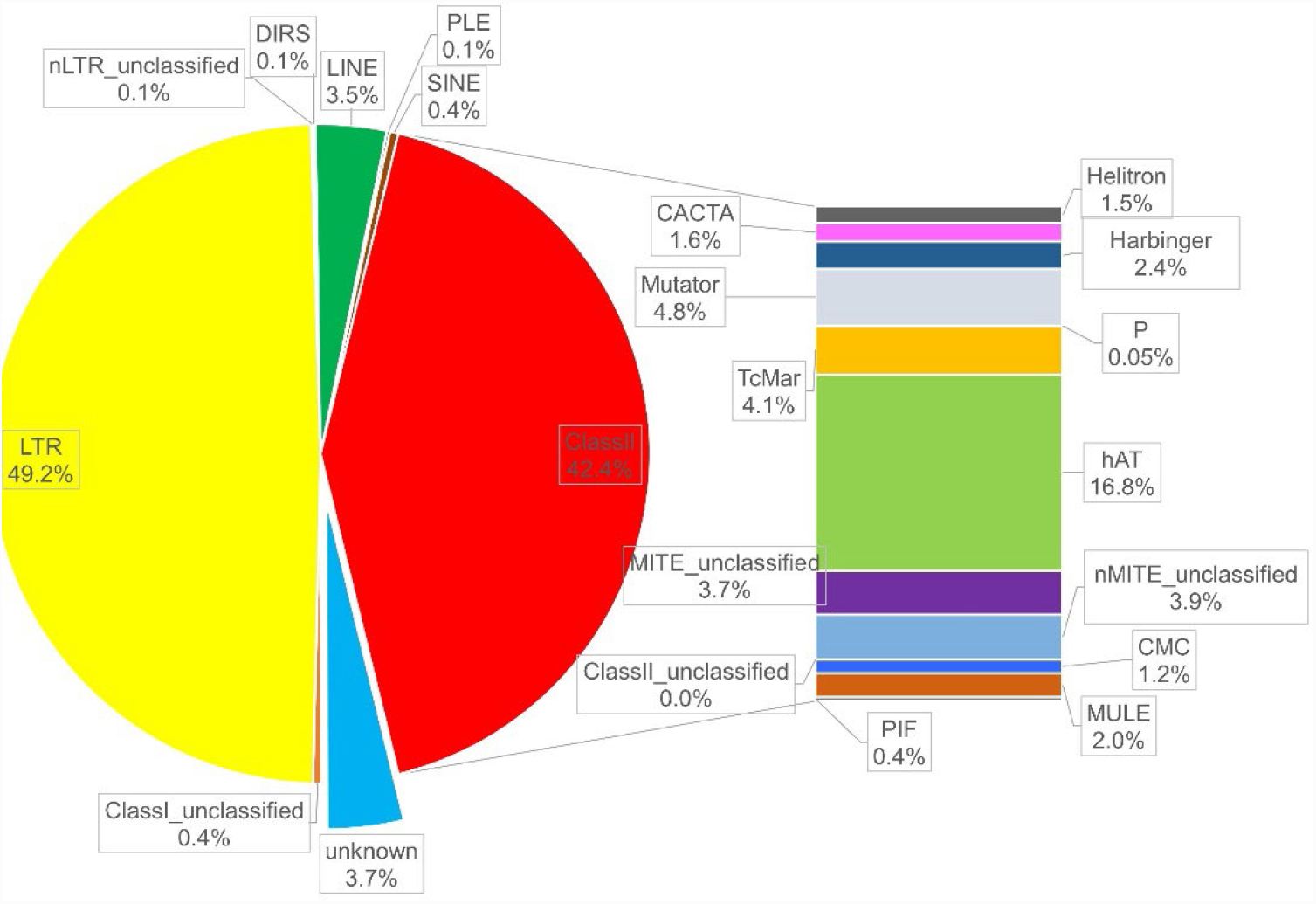
Distribution of whole-genome TE elements in different TE types. 53.9% TEs were classified as Class I (Retrotransposons), and 42.4% were classified as Class II (DNA transposons). LTRs (152.8 Mb) account for 91.3% of Class I TEs, and LINEs (10.8 Mb) are the second most abundant Class I TEs. hAT (52.3 Mb), Mutator (15.0 Mb), and TcMar (12.9 Mb) are the most abundant Class II TE types.

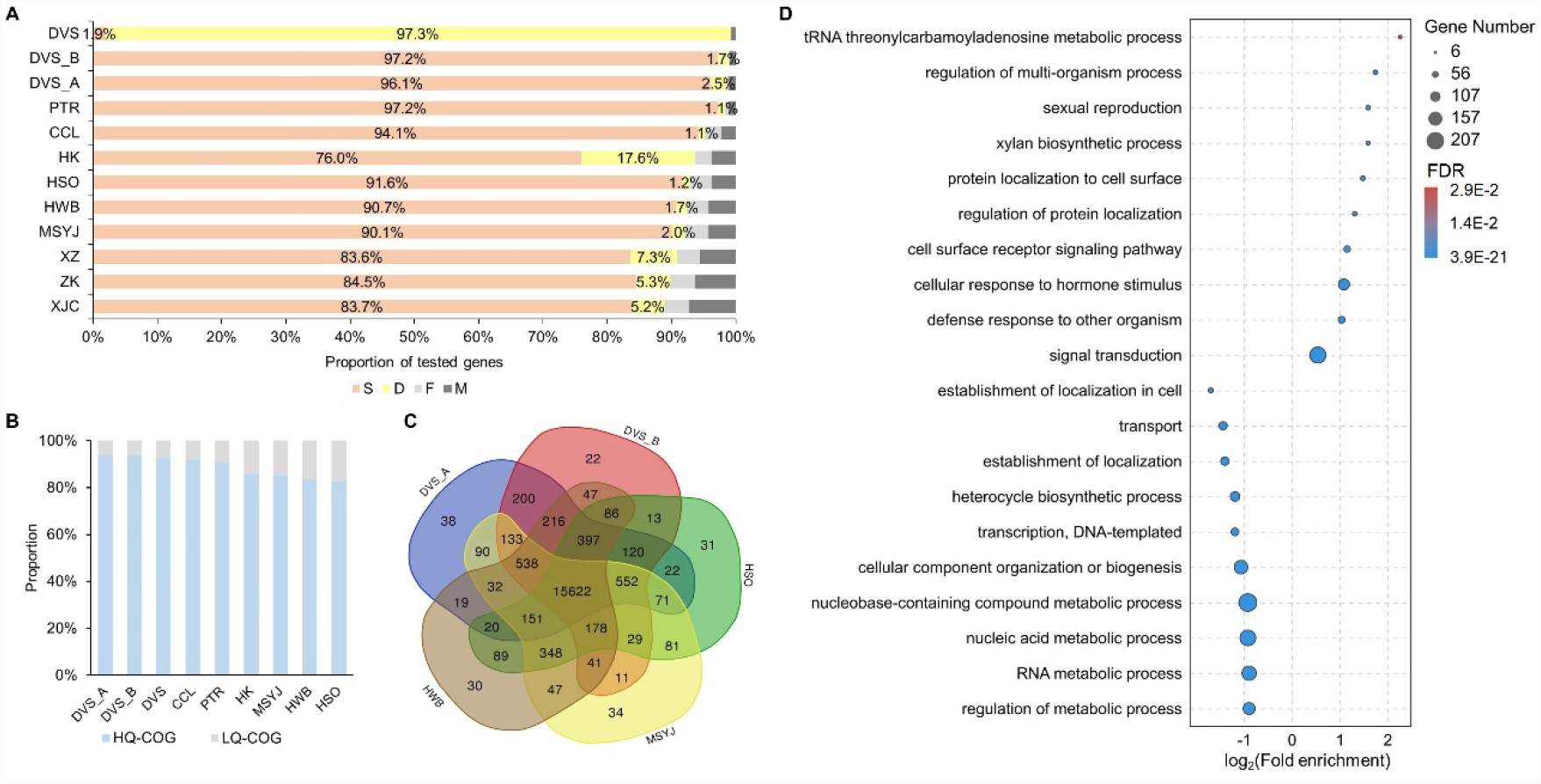

**Extended Data Fig. 5:**
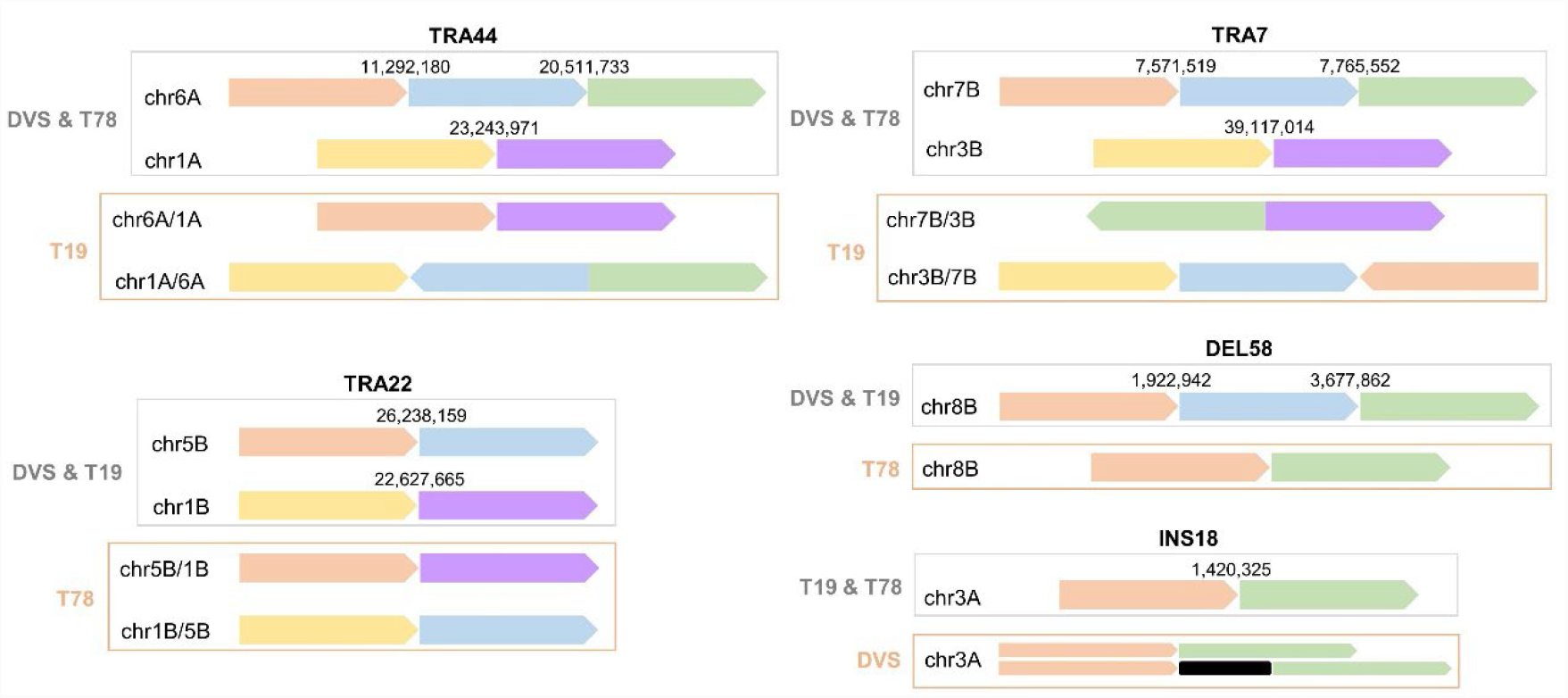
Diagrams of five somatic structural mutations detected in T19, T78, and DVS. The arrows denote the directions from 5’ to 3’ on the DVS chromosomes. The gray font and frame indicate the ‘wild type’ genotype, and the light red font and frame indicate the mutant accession and genotype of the corresponding mutation, respectively. The black bar in DVS of INS18 represents the 6,924 bp Mutator transposon. The wild type and the mutant type chr3A coexist with approximately 1:1 ratio in the chimeric DVS on INS18.

**Extended Data Fig. 6:**
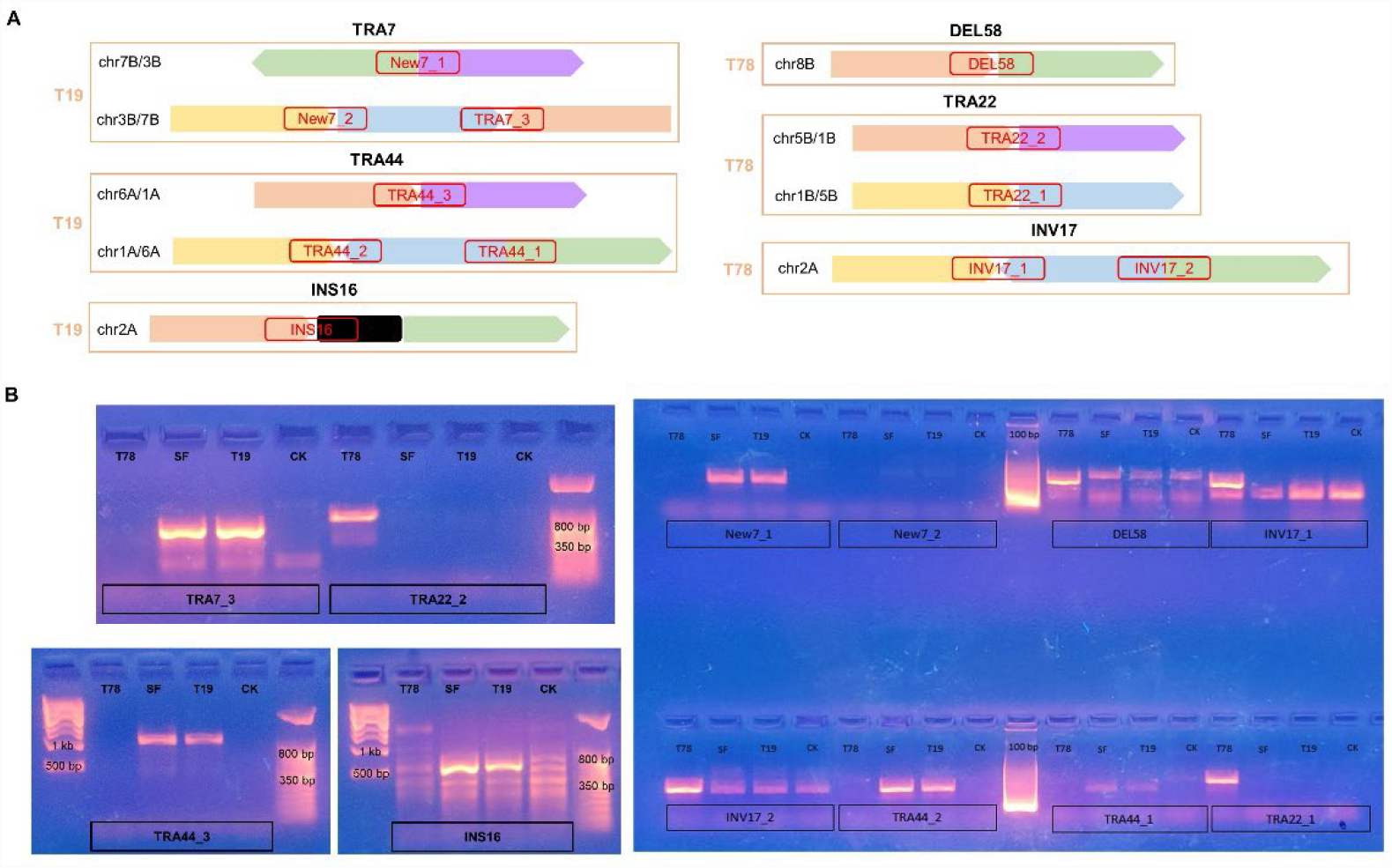
PCR verification of 6 putative irradiation-induced structural mutations. **A**. Diagrams showing the breaking end connections (connections between arrows or squares of different colors) in the mutant accession and the corresponding amplicons (red frames) designed to verify them. The arrows in panel A denote the directions from 5’ to 3’ on the DVS chromosomes. Only the mutant genotypes (the orange frames) and accession names are shown in the graph. **B**. PCR amplification of the designed amplicons. CK denotes the ordinary Valencia sweet orange DVS. The primers for DEL58, INV17_1, INV17_2, and INS16 had non-special amplifications in the non-mutant accessions, but the target amplicons were only amplified with high efficiency in the corresponding mutant accession(s).

**Extended Data Fig. 7:**
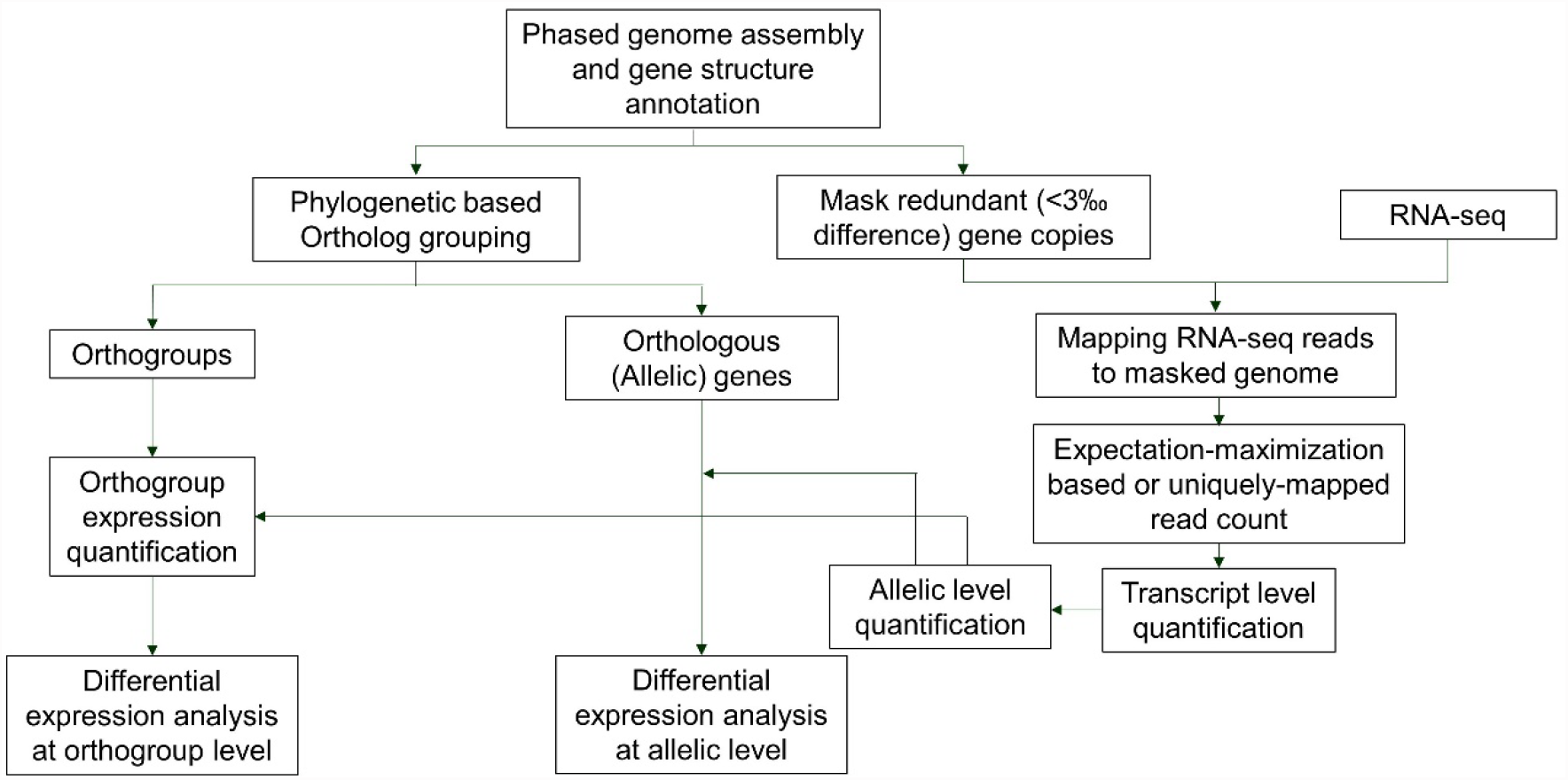
Allele-aware RNA-seq analysis pipeline for SWO.

**Extended Data Fig. 8:**
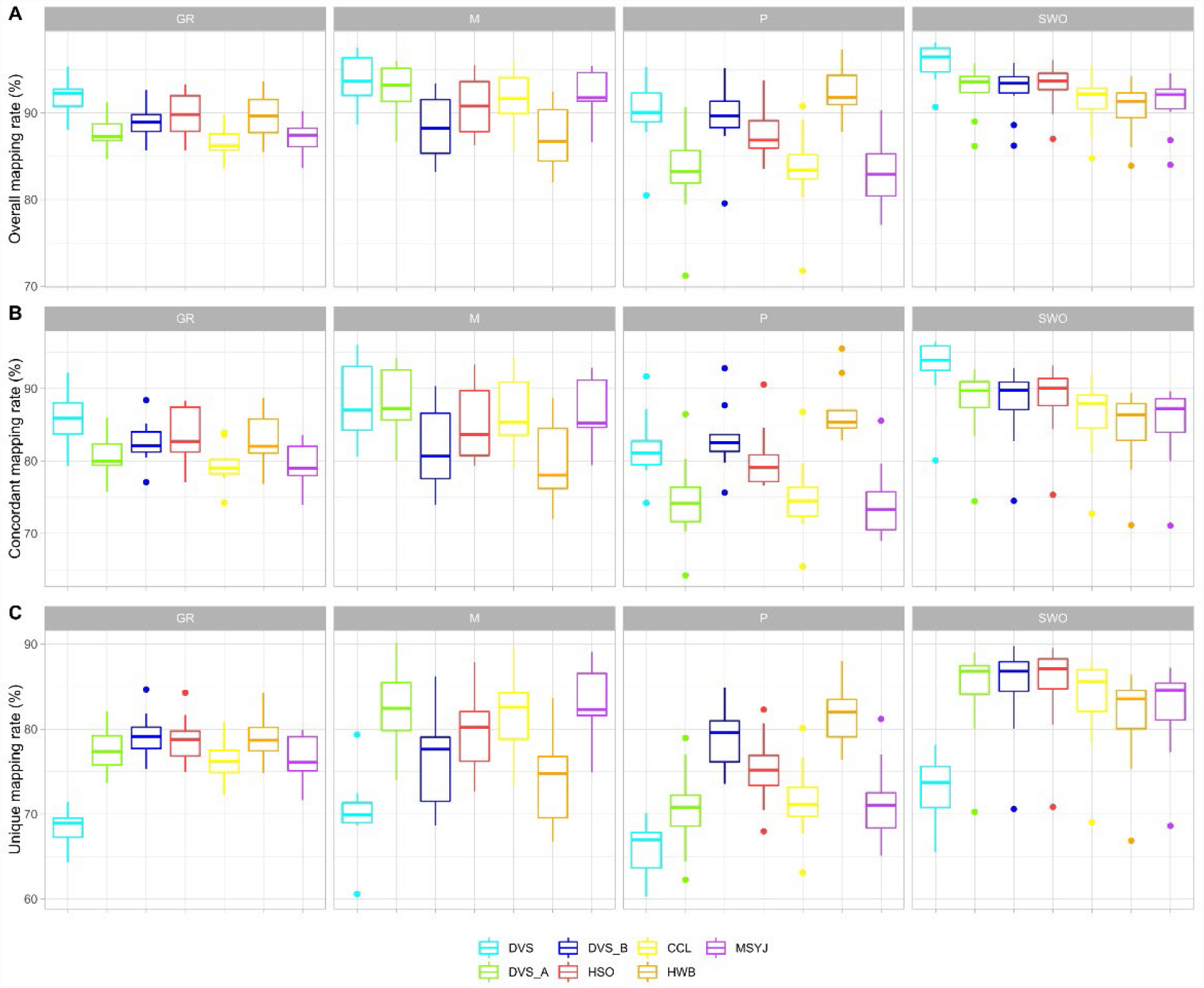
Mapping rates of RNA-seq data with different citrus assemblies as the reference. For each of the four species, grapefruit (GR), mandarin (M), pummelo (P), and SWO, ten RNA-seq data were mapped to all the tested citrus assemblies. The boxplots describe the distribution of overall mapping rates (**A**), concordant mapping rates (**B**), and the unique mapping rates (**C**). The overall, concordant, and unique mapping rates were calculated as the ratios of all mapped reads, concordantly mapped read pairs, and uniquely and concordantly mapped read pairs, respectively.

**Extended Data Fig. 9:**
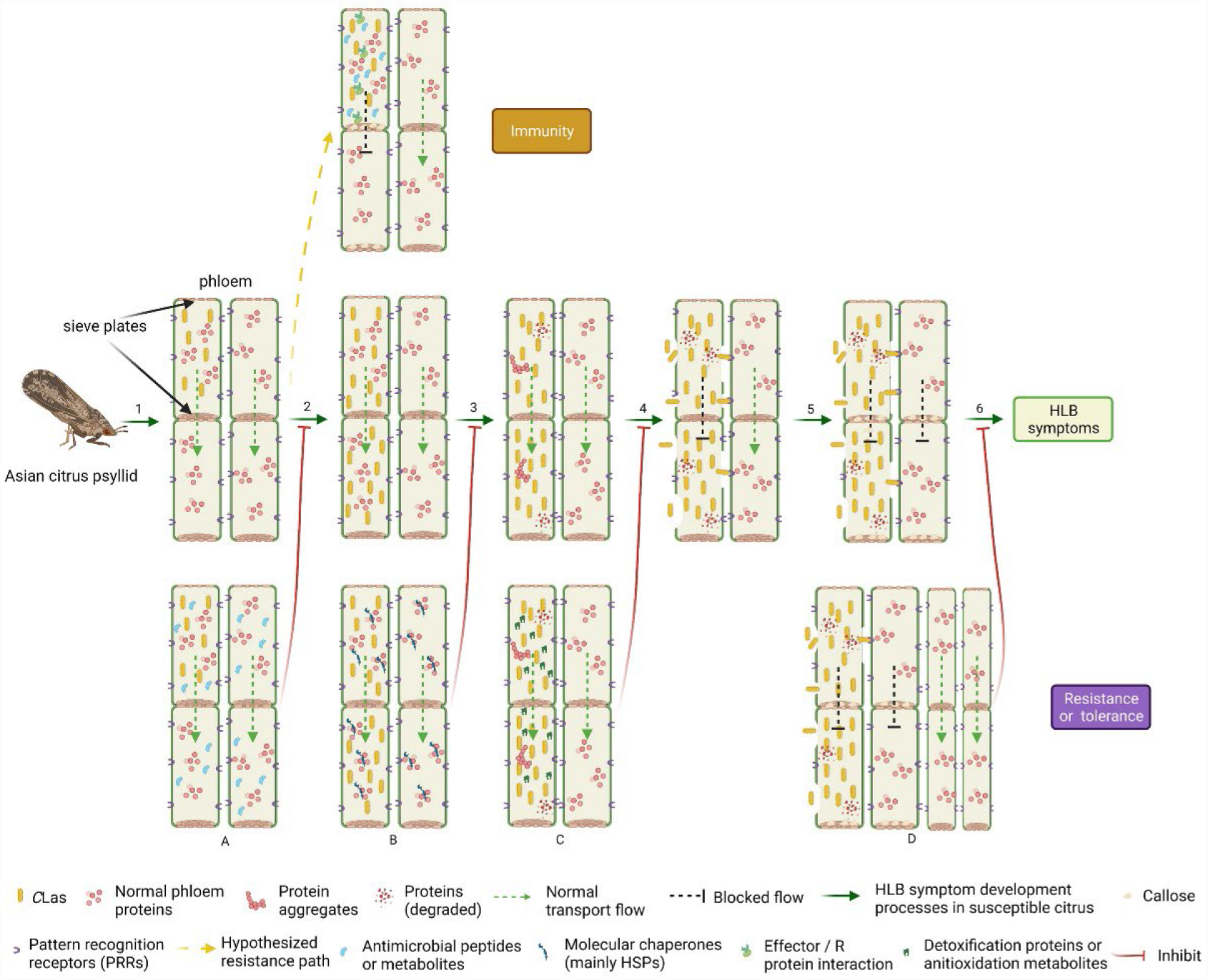
Huanglongbing (HLB) symptom development in susceptible citrus genotypes and the mechanism of HLB tolerance and resistance. Steps 1-6 denote the HLB symptom development process in susceptible citrus genotypes. 1, the pathogens are injected by Asian Citrus Psyllid (*Diaphorina citri* Kuwayama) into the phloem^8^; 2, *C*Las propagates without inducing any symptoms/(strong) defense responses for months^9^; 3, a few HSP encoding genes are significantly downregulated^46,61–63^, and reactive oxygen species (ROS) are accumulated under stress^110^, resulting in protein misfolding, aggregation, and degradation; 4, ROS and protein aggregates induce programmed cell death^60,111^ and release *C*Las from the phloem; 5, microbe-associated molecular patterns of *C*Las are recognized by pattern recognition receptors and induce defense responses^65^, including callose deposition and programmed cell death; 6, high rate of dysfunctional phloem cause symptoms including blotchy mottles, yellow branches, and eventual tree dieback^8^. A, B, C, and D are hypothesized mechanisms underlying enhanced HLB tolerance or resistance in citrus. A. Enhanced NPR1 dependent defense^41,42^ or high-level antimicrobial peptide expression^43–45^; B. Promoted protein and cellular homeostasis due to upregulation of molecular chaperones (mainly heat shock proteins); C. Inhibition of programmed cell death by high-level of antioxidant metabolites or enhanced detoxication pathways^46,48,64^; D. Increased functional phloem ratio from novel high phloem regeneration rate^15,49^.

